# BOLD fMRI signals of visual white matter encode visuotopic information and predict effective connectivity between visual areas

**DOI:** 10.1101/2021.06.30.450520

**Authors:** Huan Wang, Xiaoxiao Wang, Yanming Wang, Du Zhang, Yifeng Zhou, Bensheng Qiu, Peng Zhang

**Author notes:** **Correspondence:** Peng Zhang, Bensheng Qiu, Yifeng Zhou. These authors contribute equally to the work (Huan Wang, Xiaoxiao Wang).

## Abstract

The functional significance of BOLD signals in white matter (WM) remains unclear. The current study investigated whether 7T BOLD fMRI signal from visual WM tracts contains high fidelity retinotopic information and whether it correlates with the effective connectivity between visual areas. Population receptive field (pRF) analysis of the 7T retinotopy dataset from the Human Connectome Project revealed clear contralateral retinotopic representations from two visual WM bundles: optic radiation (OR) and vertical occipital fasciculus (VOF). The retinotopic organization of OR is consistent with post-mortem studies. The pRF size of WM voxels also increases with eccentricity. Based on the retinotopic maps of OR, we investigated whether BOLD signals in OR during visual stimulation are related to the resting-state effective connectivity between the lateral geniculate nucleus (LGN) and the primary visual cortex (V1). Results show that visually-evoked BOLD responses in OR correlate with the feedforward and feedback connectivity between the LGN and V1 during resting state. These findings demonstrate that WM BOLD signals contain high fidelity information such as visual field maps, and also predict the functional connectivity of brain areas.

**Significance statement:** White matter (WM) tracks conduct spiking activity between distant neurons. Weak fluctuations of BOLD signals in the WM can be detected with fMRI, but their functional relevance remains largely unknown. Here we characterized the visual field map properties of two major visual WM bundles: the optic radiation (OR) and vertical occipital fasciculus (VOF). Population receptive field analysis of the WM BOLD signals revealed clear visual field maps in both WM tracts. Effective connectivity analysis further showed that visually evoked BOLD responses in OR can predict the resting thalamo-cortical functional connectivity. These findings demonstrate that WM BOLD signals contain highly specific functional information and could directly index the functional connectivity between brain areas.

## Introduction

White matter (WM) is composed of bundles of axons transferring spiking activity between distant neurons in the gray matter (GM). A non-invasively tool to directly access the functional activity of WM is crucial for understanding the information transfer of the human brain. It is well recognized that, due to rich vasculature and neurovascular coupling, neural activity in the GM lead to blood-oxygen-level-dependent (BOLD) activity that can be sensitively detected by functional magnetic resonance imaging (fMRI). BOLD signals in the WM are much weaker due to much less dense vasculature. Study on GM has found that fMRI activations extent well beyond the areas of primary relation to the task, under optimal noise conditions (1). It is likely that WM also exists this phenomena of similar nature even with increased noise level (2). Recent findings show that WM BOLD signals are still measurable with fMRI (3-5), but its functional significance remains an open question.

Several fMRI studies detected BOLD signals from the WM during task or sensory stimulations (5-7), modulated by task load or stimulus parameters in similar ways as the GM areas they connected (2). During resting state, BOLD fluctuations in the WM also showed similar spectral profiles as in the GM and can be used to reveal the functional connectome of the brain (5, 8-10). These findings suggest that WM BOLD signals could be related to the intrinsic neural activity. However, whether WM BOLD signals contain highly functionally specific information, and its relationship with the functional connectivity of the brain require further investigation. Neurons in the visual areas of the brain tune to spatial locations of visual stimuli in their receptive fields. Population receptive field (pRF) mapping on BOLD fMRI signals of GM has successfully revealed visual field maps in many cortical and subcortical areas (11-13). In the current study, we used the pRF method to investigate whether high fidelity visuotopic information can be derived from the BOLD signals of visual WM tracts. Dynamic casual modeling (DCM) was also used to investigate whether WM BOLD activity correlates with the effective connectivity of brain areas.

Optic radiation (OR) is the WM tract transferring information between the lateral geniculate nucleus (LGN) of the thalamus and the primary visual cortex (V1). Lesions to OR lead to blindness of the corresponding visual field (14). The retinotopic organization of OR was previously investigated by post-mortem and diffusion MRI studies (15, 16). Abnormality in micro-structure integrity of OR is closely linked with many ocular and brain diseases, such as amblyopia, glaucoma and schizophrenia (17-19). A non-invasive method to evaluate the function of OR should be highly valuable to study the neural mechanism of these diseases. Vertical occipital fasciculus (VOF) is the fiber bundle connecting dorsolateral and ventrolateral visual cortex (20, 21). It plays critical roles in processing higher level visual information, such as stereopsis, object category information, and visual-motor control signals (20-22). To the best of our knowledge, the retinotopic organization of VOF has not been investigated. Therefore, if BOLD fMRI can reveal functionally specific retinotopic activity of visual WM tracts, it can be useful to study the information transfer of the brain in many neuroscience and clinical applications.

Since the sensitivity and specificity of BOLD fMRI signals increase with the magnetic field strength (23), we used the 7T retinotopic dataset from the Human Connectome Project (HCP) (24) to characterize the visual field map properties of OR and VOF. The pRF analysis revealed robust polar angle and eccentricity maps for both visual WM bundles, and the map of pRF size is highly consistent with the eccentricity map. Estimated hemodynamic response functions (HRF) of both WM tracts are significantly delayed compared to the canonical HRF of the visual cortex. DCM analysis showed that OR BOLD activity to visual stimulation significantly correlated with the thalamocortical effective connectivity between the LGN and V1 during resting-state.

## Results

### pRF estimates of OR and VOF

To estimate the visual field map properties of the two visual WM bundles, we employed the preprocessed 7T retinotopy datasets from HCP, at both high spatial (1.6mm isotropic) and high temporal (1000 ms) resolution (24). ROIs for the OR and VOF bundles were traced from the group-averaged diffusion images using a deterministic fiber tracking algorithm (25) through DSI Studio (http://dsi-studio.labsolver.org) (26). Figure 1A&C show the tractographies. Retinotopic fMRI data from all subjects were normalized to Montreal Neurological Institute (MNI) space with the ICBM2009c T1 template (27, 28). Voxel-wise pRF parameters were estimated from the group-averaged fMRI time series within the ROIs of OR and VOF. Example time courses in the right panels of Figure 1B&D show that BOLD responses in OR and VOF can be fitted reasonably well by the pRF model.

**Figure 1.**
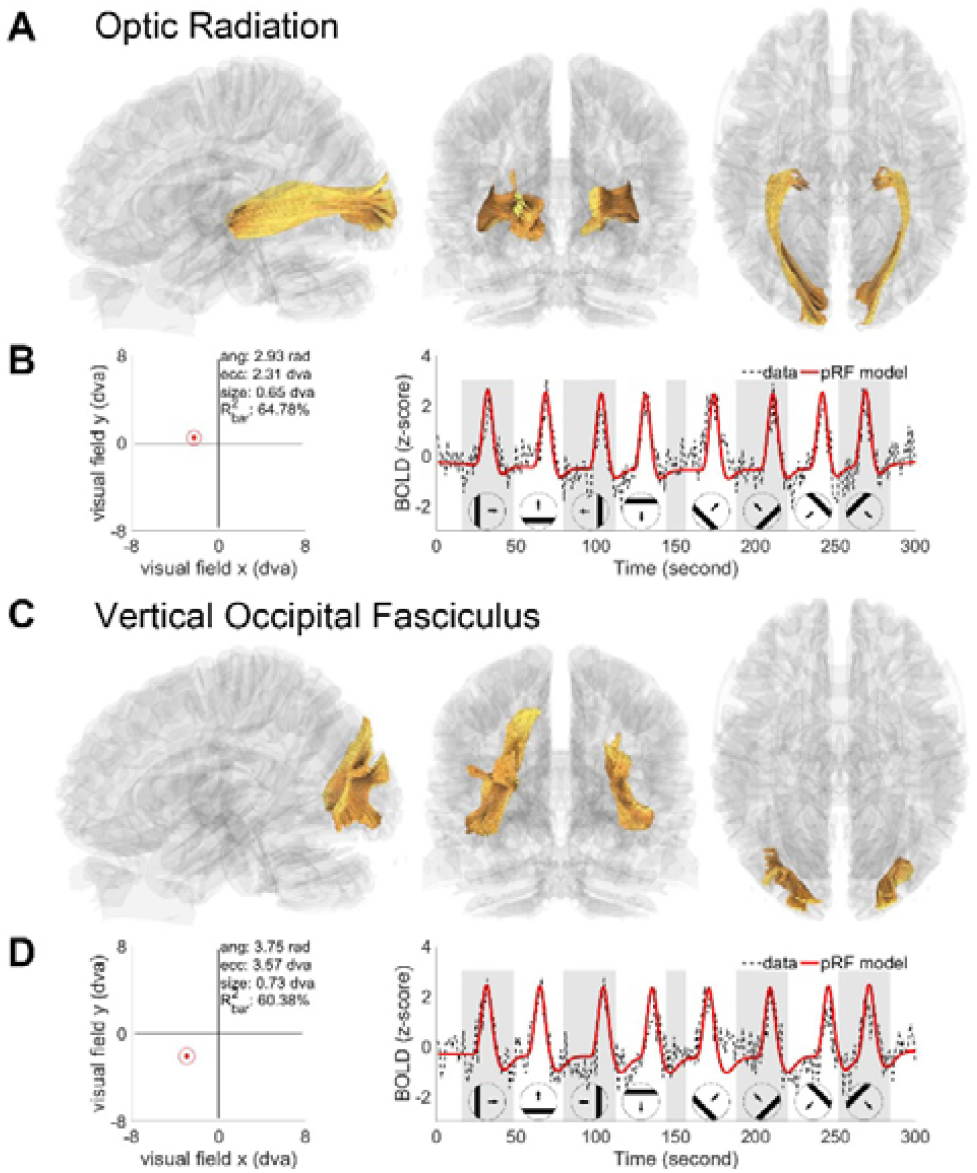
WM fiber bundles and pRF fitting examples from the group-averaged data. (A) Sagittal, coronal and axial views of tracked OR fiber bundle. (B) shows the pRF profile estimated from an example OR voxel, the red dot represents the pRF center and the red circle represents the scope of pRF. The right panel shows the coupling between example time courses (average across two bar stimuli runs, black dashed line) and pRF model predictions (red solid line). R^2^_bar_ indicates the variance explained by the pRF estimates. (C) Tractography for VOF fiber bundle. (D) pRF estimates for an example VOF voxel. Original and fitted fMRI responses are shown on the right. dva, degrees of visual angle.

The retinotopic maps of OR are illustrated in coronal sections in Figure 2. The polar angle maps show clear contralateral representations. For each hemifield, the upper and lower visual field are represented at the ventral and dorsal portion of OR, respectively. Voxels representing the central visual field locate at the middle portion of OR, while peripheral representations locate more dorsally and ventrally. These retinotopic organizations are consistent with previous post-mortem and DTI studies (15, 16). The posterior slices show higher explained variance and more clearly represented visual field maps, likely due to higher signal-to-noise-ratio of 7T fMRI signals in the occipital cortex than in the middle of the brain. Most OR voxels have small pRF sizes, with larger pRFs at larger eccentricities. The map for the time-to-peak (TTP) of HRFs shows a relatively uniform distribution in OR. Figure S1 shows the pRF mapping results from a representative subject.

**Figure 2.**
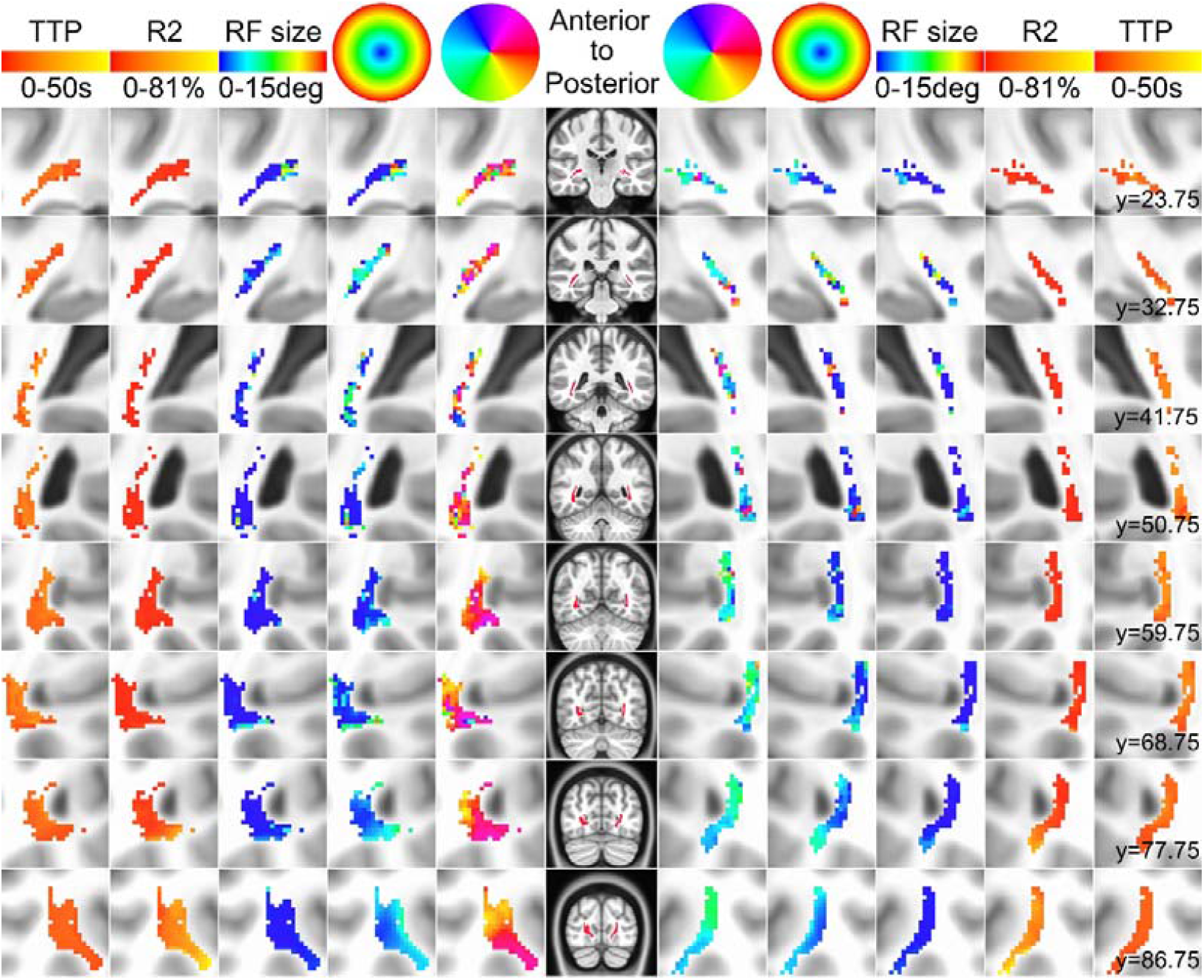
Estimated visual field maps and HRF delays for OR. Results are presented in selected coronal slices. Voxels with 0.68% of variance explained were shown. Columns from middle to left and right sides illustrate the OR mask, polar angle, eccentricity, pRF size, explained variance and TTP of HRF, respectively. Rows represent separate slices, arranged from anterior to posterior. MNI coordinate of y axes are shown in millimeter.

The pRF analysis also revealed robust contralateral representations of visual field from the BOLD signals of VOF connecting the ventral and dorsal visual areas (Figure 3). The polar angle and eccentricity representations are mirror symmetric between the two hemispheres and smoothly distributed, i.e. nearby voxels represented nearby visual fields. Unlike OR, the VOF seems to represent multiple lower-upper visual field representations within each axial slice. The eccentricity maps are over-represented in the central visual field, consistent with the magnification factors of the LGN and V1. The maps for pRF size show similar orgnizations as the eccentricity maps.

**Figure 3.**
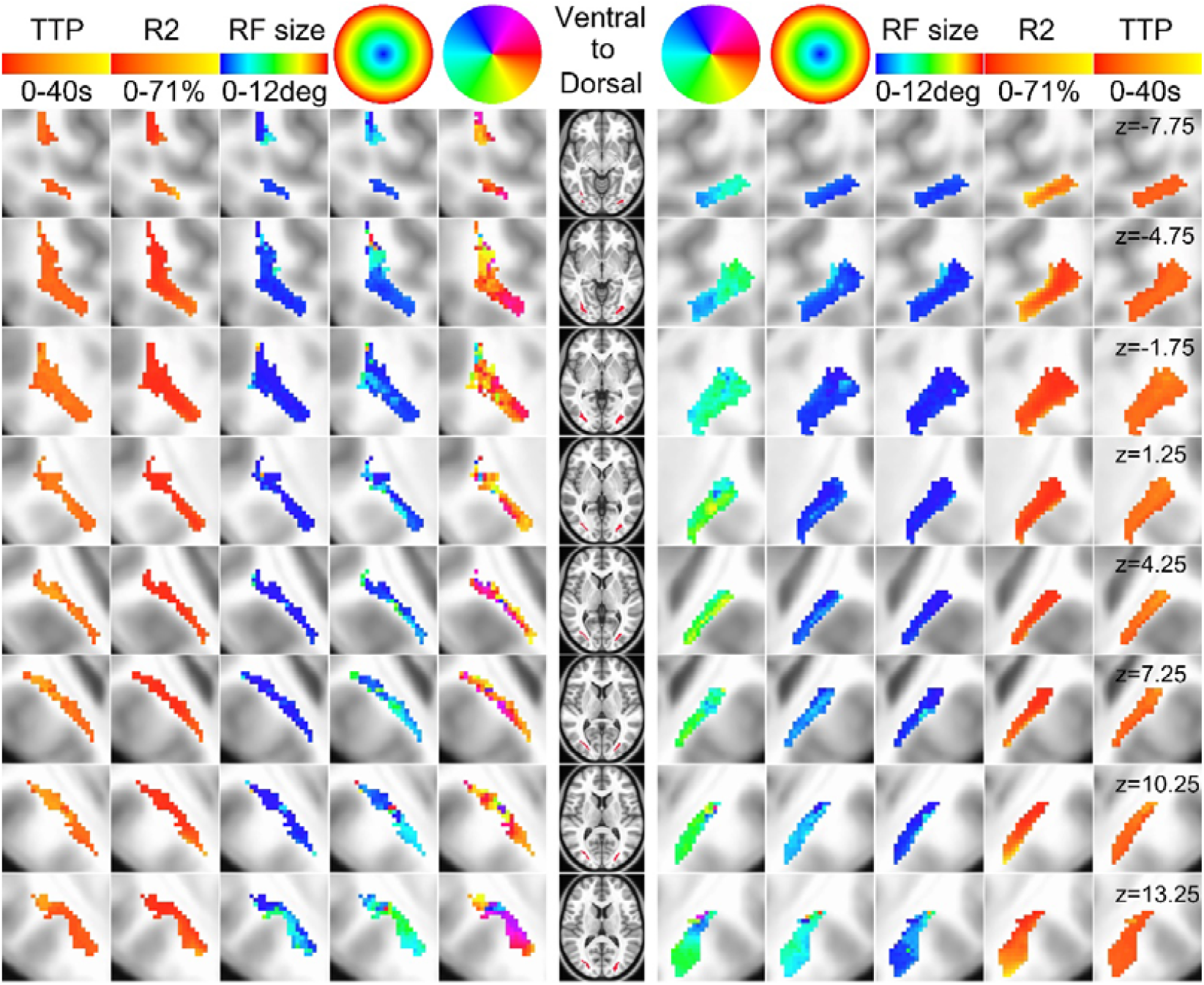
Estimated visual field maps and HRF delay in VOF. Selected axial slices arranged from ventral to dorsal visual cortex. Results were thresholded at 0.31% of variance explained. Conventions as in Figure 2.

For both visual WM tracts, the polar angle representations show strong bias around the horizontal meridian, with much less polar angle representations around the vertical meridian (Figure 4A&E). Moreover, the representation of VOF is strongly biased towards the lower visual field. The over-representation of visual field around the horizontal meridian is consistent with a previous retinotopic fMRI study of the human LGN (29). Eccentricity distributions show strong foveal (eccentricity < 4 degree) bias for both OR (79.38% of the voxels, Figure 4B) and VOF (79.64% of the voxels, Figure 4F). The size of pRF also increases with eccentricity, as indicated by significant positive correlations between the two parameters in figure 4C&G (r=0.62, p<0.0001 for OR; r=0.51, p<0.0001 for VOF). A double-gamma function for the HRF was fitted together with the pRF model parameters. Figure 4D&H show the fitted HRFs for OR and VOF. Compared to the HRFs of early visual cortex with a time-to-peak (TTP) at about 4-5 s (30), we found much delayed HRFs for both visual WM tracts (TTP = 9.96 (mean) ± 4.47 (st.d.) s and 8.12 ± 3.94 s for OR and VOF, respectively). Figure S2 shows the results from a representative subject. This finding is consistent with a recent study showing delayed WM HRFs in fronto-parietal regions (31). To summarize, results from the pRF analysis provide strong evidence that WM BOLD signals contain highly specific functional information such as visual field maps.

**Figure 4.**
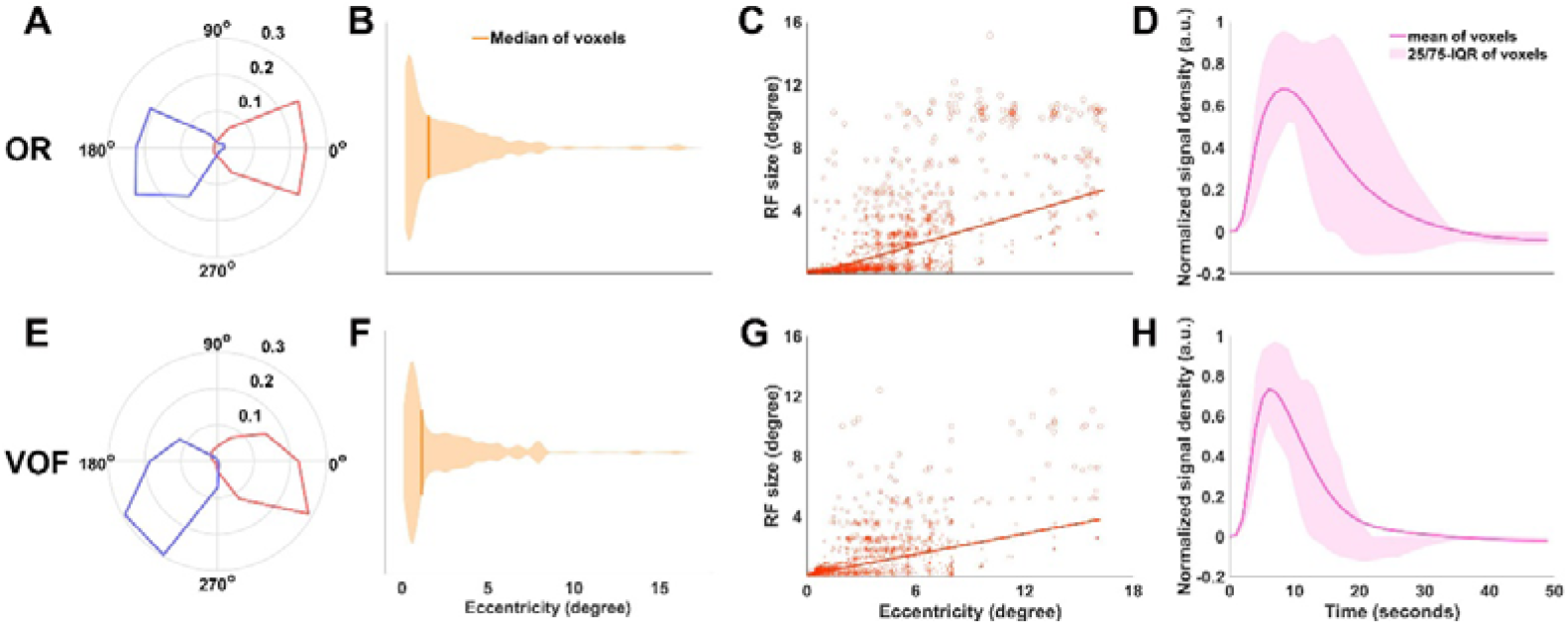
Model parameters of OR (A, B, C, D) and VOF (E, F, G, H) for group average level (across 178 HCP subjects). Survived voxels after thresholding (at 0.68% of variance explained for OR and 0.31% of variance explained for VOF) are shown. (A&E) Polar angle distribution. The solid lines stand for the fractional volume representing each polar angle. The left (red) and right (blue) hemispheres are plotted separately. (B&F) Eccentricity. More than 75.69% OR voxels and 75.61% VOF voxels represent central 4 degree visual field. (C&G) pRF size versus eccentricity. The solid lines are linear fits and gray points stand for each voxels. (D&H) Double gamma HRF model fitted HRF curve for OR and VOF. HRF curves of each voxel were normalized to their peak magnitude. IQR is interquartile range.

### Visually evoked BOLD activity in OR correlates with the effective connectivity between the LGN and V1 during resting-state

Axons in the WM transfer information between brain areas via spike trains. If WM BOLD signals indeed reflect the neural activity of axons, the BOLD responses in the WM should be related to the information transmission or functional connectivity of connected brain areas. However, this intuitive prediction has not yet been demonstrated. To test this hypothesis, we investigated the relationship between visually evoked OR BOLD activity and the resting-state effective connectivity between the LGN and V1. Dynamic casual modelling (DCM) (32) was used to derive the effective connectivity between the two visual areas, followed by a second-level parametric empirical Bayes (PEB) analysis (33, 34) with the visually evoked OR BOLD responses as a covariate of between-subject effect. Due to the potential difference in feedforward and feedback activities for central and peripheral vision (35, 36), OR and both visual areas were separated into central and peripheral ROIs using the visual field maps generated from the pRF analysis. The DCM-PEB analysis was performed separately on the central and peripheral ROIs.

In figure 5A, the fixed connections of DCM model includes both feedforward (FF) and feedback (FB) connections between the LGN and V1, as well as their self-connections. The full model for PEB assumes that visually evoked OR BOLD activity can correlate with both FF and FB connections. Reduced models define either FF or FB connection for the effects of OR BOLD activity. Figure 5B-D show the results for the central visual field. Bayesian model comparisons reveal the best model as FF/FB (with the highest exceedance probability at 68%). After Bayesian model average (BMA), parameter estimate for the fixed connections show both significantly positive FF and negative FB connections, as well as negative self-connections for both visual areas. The effect of OR BOLD activity indicates a significant positive correlation with the FB connection from V1 to the LGN (posterior probability (Pp) = 100%). For the peripheral visual field, the FF model shows the highest exceedance probabilities for the effect of OR BOLD activity (64%) (Figure 5E). The FF connection parameter related to the OR BOLD activity also shows high posterior probability (Pp = 89%) (Figure 5F). To summarize, the DCM-PEB analysis shows that visually evoked OR BOLD activity is closely related to the functional connectivity between the LGN and V1 during resting-state.

**Figure 5.**
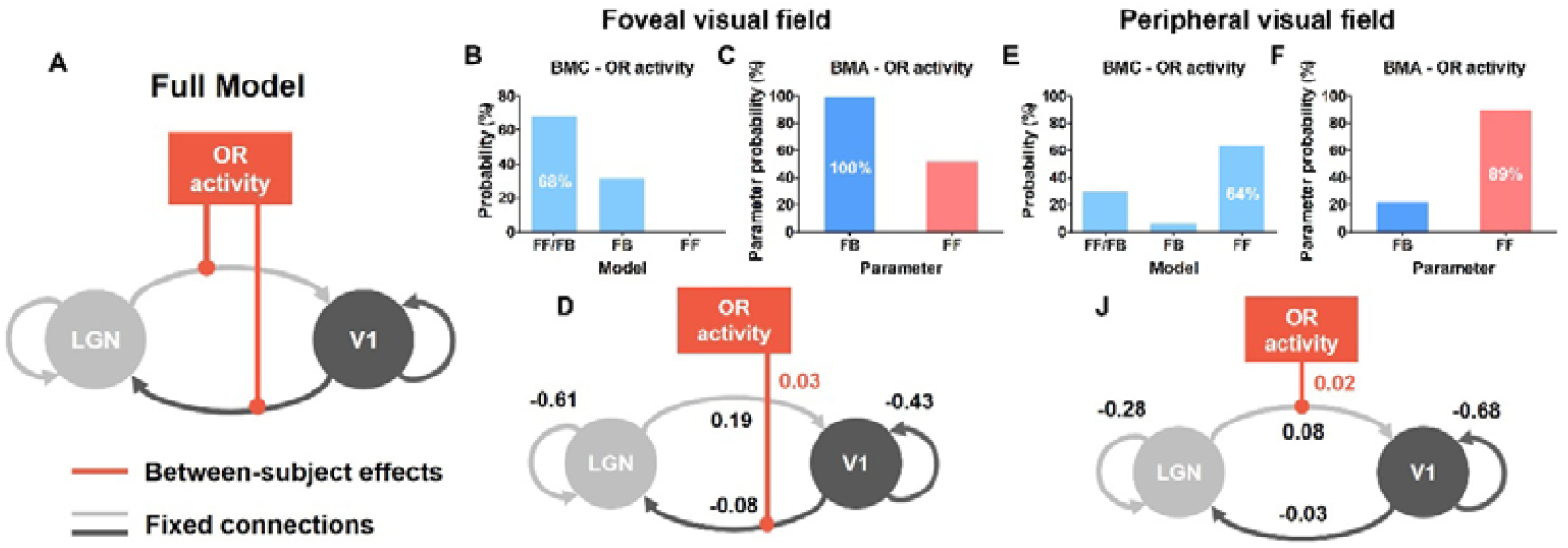
Effective connectivity analysis model and results. (A) The full DCM-PEB model included self-connections and feedforward (FF) and feedback (FB) connections between LGN and V1, and with between-subject effect of OR BOLD activity on both feedforward and feedback connections (red solid lines). (B) Model exceedance probabilities for the effect of OR BOLD activity in the central visual field. The full model FF/FB has 68% posterior probability for OR activity. (C) Posterior probability of connection parameters in the central visual field. (D) Parameter estimates for the central visual field. OR BOLD activity showed a significant positive effect on the FB connection from V1 to LGN (effect size = 0.03, with about 100% posterior probability). (E) Model exceedance probabilities in the peripheral visual field. (F) Posterior probability of connection parameters in the peripheral visual field. (J) Parameter estimates for the peripheral visual field.

## Discussion

Using pRF analysis on the WM BOLD fMRI data from the 7T retinotopic dataset of HCP, we characterized the visual field map properties of two major visual WM bundles, OR and VOF. Clear polar angle and eccentricity representations were found for both WM tracts. Eccentricity maps reveal over-representations of the central visual field, with the pRF size increased with eccentricity. The VOF also shows a strong bias towards the lower visual field. The retinotopic organizations of OR are consistent with previous post-mortem and DTI studies (15, 16). For the first time, we revealed the retinotopic organization of VOF. HRFs of both visual WM tracts show significantly delayed time-to-peak response compared to the canonical HRF from the visual cortex. Finally, the DCM-PEB analysis indicates that visually evoked OR BOLD activity is closely related to the effective connectivity between the LGN and V1 during resting-state.

Although BOLD signals of WM have been detected by fMRI in previous studies (5-8), its functional relevance remains elusive (37). The current study provides strong evidence that WM BOLD signals contain highly functionally specific information such as visual field maps, suggesting that BOLD fMRI can be used to evaluate the function of WM in a fiber-specific way. The ability to measure retinotopic activity of visual WM can be valuable to study the axon-related neural mechanisms in many visual and brain diseases such as glaucoma, macular degeneration, hemianopia, amblyopia and schizophrenia.

The DCM effective connectivity analysis provides direct evidence that the BOLD activity of WM is closely related to the functional connectivity of connected brain areas. Results show that visually evoked OR BOLD activity from the central and peripheral visual field correlated with the feedback and feedforward resting-state connectivity between the LGN and V1, respectively. This finding is consistent with the view of central-peripheral dichotomy that central vision requires stronger feedbacks for perceptual inference, a process that could be much weaker in the peripheral visual field (36). It is also consistent with the findings from recent studies that spontaneous connectivity patterns in the early visual areas are eccentricity-dependent (38, 39). Therefore, in addition to the current measurements of functional connectivity that contributed by both direct and indirect connections, BOLD activity in the WM could be a useful physiological measurement to assess the direct functional connectivity of connected brain areas.

One major challenge for the interpretation of WM BOLD signals is the possible contamination from adjacent GM voxels. In the current study, GM signals cannot explain the visual field maps of WM for the following reasons: 1) The observed retinotopic maps from OR BOLD signals are consistent with the organization of OR from previous studies, but not those from nearby visual cortex; 2) HRFs of both WM tracts are much delayed compared to the canonical HRF from GM, consistent with the HRF properties of WM (31); 3) A rigid mask of 90% white matter probability was generated from the results of FreeSurfer volume parcellation at the individual level.

To conclude, the current study demonstrates that high-fidelity neural information such as visual field maps can be derived from the BOLD signals of visual white matter. Visually evoked BOLD activity of WM can also predict the effective connectivity of connected visual areas during resting-state. These findings significantly improved the current understanding about the functional relevance of WM BOLD signals. The visual field maps for OR and VOF could be useful to study the axon-related neural mechanisms in many neuroscience and clinical applications.

## Methods

### Subjects

The current study contained 178 Young Adults from the Human Connectome Project (109 females, 69 males), age from 22 to 35. All subjects have normal or correct-to-normal visual acuity. Each subject has an assigned six-digit HCP ID.

### HCP neuroimaging data acquisition

We downloaded 178 subjects’ datasets from the HCP database (https://db.humanconnectome.org). The 3T datasets include two packages: (1) Structural Preprocessed, which contained 0.7mm isotropic spatial resolution structural data in standard MNI space preprocessed with the HCP structural pipeline; (2) Structural Extended Preprocessed, which was surface-based data by the HCP structural pipeline, was used for defining V1 ROIs for DCM modelling. The 7T datasets we used included four packages: (1) Resting State fMRI Functional Preprocessed Extended, volume format data with 1.6mm isotropic spatial resolution in standard MNI space; (2) Retinotopy Task fMRI Functional Preprocessed Extended, which was furtherly normalized to ICBM152 2009c space through the nonlinear warp from the Structural Preprocessed data to the 1.0mm isotropic ICBM152 2009c template, with the help of the ANTs software (40); (3) Structural Preprocessed for 7T, structural data has the same resolution and space with fMRI data; (4) Diffusion Preprocessed, 1.05mm spatial resolution diffusion and structural data in the native MNI space. TR for both fMRI datasets is 1000ms. HCP pipelines for data collection and preprocessing are extensively described in Benson, *et al*. (24), Glasser, *et al*. (41), A, *et al*. (42).

### Stimuli for retinotopy dataset

The stimuli used for generating retinotopy datasets were dynamic colorful textures in the slow-moving apertures, including clockwise or counter-clockwise rotating wedge (RETCW, RETCCW), expanding or contracting ring (RETEXP, RETCON) and moving bar (RETBAR1, RETBAR2). Stimuli were generated in the resolution of 768*768 pixels and were constrained to a circular region with the diameter of 16.0 degrees of visual angle. The order of stimuli was RETCCW, RETCW, RETEXP, RETCON, RETBAR1, and RETBAR2. These stimuli corresponded to 6 scan runs of 5 minutes long (300 fMRI time points) each. Other details about stimuli can be found in the literature of Benson, et al. (24).

### Fiber tracking

Fiber tracking analysis was conducted using the DSI Studio (http://dsi-studio.labsolver.org/). A deterministic fiber tracking algorithm (25) was performed to identify the optic radiation (OR) and the vertical occipital fasciculus (VOF) for pRF estimating of group average level. The 1-mm spin distribution function (SDF) population-averaged template HCP-1065 (ICBM152 2009a space) (26) provided by DSI Studio for tracking visual fiber bundles was used.

Angular thresholds of 60, 70 and 80 degree were used for conducting the fiber tracking from the LGN to the V1. Each angular threshold generated 20000 streamlines, which were then clustered using single-linkage clustering (26). Clusters with less than 200 streamlines were excluded. Then the remaining streamlines from each angular threshold were merged and the ROI of OR was got. The VOF fiber pathway was described as a major white matter fascicle connecting dorsal and ventral visual cortex by the previous literatures (21, 43). Therefore, the VOF fiber pathway were tracked between regions of hV4/VO-1 in ventral and V3A/B in dorsal, with angular thresholds of 40, 50 and 60 degree. The next processes were similar as the OR fiber tracking. The ROI of lateral geniculate nucleus (LGN) was drawn manually on MNI ICBM 2009c Nonlinear Asymmetric T1w image, according to the PDw anatomical template(13, 27, 28). The ROIs of V1, hV4/VO-1 and V3A/B were obtained from the maximum probability atlas by Wang, Mruczek, Arcaro and Kastner (44).

The OR and VOF pathways of every subject were also tracked using a batch automatic fiber tracking method (45). Each subject’s 1.05 mm isotropic SDF was obtained by the reconstruction of the diffusion MRI data with the Generalized q-sampling imaging (GQI) method (46). Then, automatic fiber tracking with default parameters was run with the templates of OR and VOF from the tractography atlas (26).

### WM masks registration and constraint

Whole brain WM mask of each subject was obtained from the wmparc results of the FreeSurfer volume parcellation (thresholded at 2600). The WM fiber tracts were converted into volumetric WM masks. For the group average level analysis, the generated OR and VOF masks were in the nonlinear ICBM152 2009a space, so they were furtherly transformed into the ICBM152 2009c space through a warp from the ICBM152 2009a template to the ICBM 2009c template by using the ANTs. To avoid signal contributions from adjacent GM voxels, a rigid group-level whole-brain WM mask was generated: First, each subject’s whole brain WM mask was warped to the ICBM152 2009c space; Then the group-level whole-brain WM mask was defined as the areas having >90% overlaps of all these warped individual WM masks. Rigid OR and VOF masks were generated by the intersection of the OR and VOF masks and the rigid group-level whole-brain WM mask.

At the individual level, each subject’s OR and VOF masks were cropped within their corresponding whole brain WM mask and then nonlinearly warped to T1w images (1.6mm spatial resolution) in the package of Structural Preprocessed for 7T.

### pRF analysis

The analyzePRF toolbox (24), based on the Compressive Spatial Summation model (47), was used to calculate pRFs of the voxels within the OR and VOF. The toolbox and the necessary codes for the HCP 7T retinotopy dataset were downloaded from the Open Science Framework web site (https://osf.io/bw9ec/). The pRF model predicts the fMRI time series as the sum of baseline signals and stimuli related signals, r(t): r(t) = (*g* × (*S*(*t*) · *G*)^*n*^ * *h*, where *g* is the gain factor, *S*(*t*) is the stimulus at time t, *G* is a 2-D isotropic Gaussian shaped receptive field, n is the power-law nonlinearity factor (fixed at 0.05 in the present study), and *h* is a parametric HRF. The pRF parameters (polar angle, eccentricity and pRF size) and HRF parameters (see below section about HRF estimation) were calculated from the fitted parameters of *G* and *h*. Please refer to the Benson, et al. (24) for more details of pRF model and fitting.

In the current study, the pRF analysis were applied both in group average data and in the individual’s data. The group average data were obtained by an average of 178 subjects’ retinotopy data, which were already warped to the ICBM152 2009c space. The pRF maps were then thresholded according to the variance explained: voxels with the variance explained lower than the threshold likely reflect noise and were abandoned. The threshold was determined by fitting a Gaussian mixture model to the distribution of the variance explained (24, 48).

### Voxel-wise WM HRF estimation

In order to investigate the delay of HRF in visual-related WM bundles, the double gamma HRF model used in the SPM (https://www.fil.ion.ucl.ac.uk/spm/) was employed in the present study:

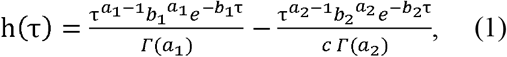

where τ is the time from the neural event, and *Γ* is the gamma function. All the five parameters (*a*_l_, *b*_l_, *c, a*_2_ & *b*_2_) were estimated during each fitting of the pRF model, and the time-to-peak (TTP) was computed from the estimated HRF curves.

### Dynamic causal modeling

176 out of 178 subjects’ resting state data (the first run) were used in DCM modelling. The eccentricity maps of LGN, V1 and OR were generated at first. The eccentricity maps of the LGN were obtained from the HCP subcortical pRF results (24). We resampled the HCP subcortical pRF results into 1.6mm spatial resolution, and then warped a careful hand-draw anatomical LGN mask in the MNI ICBM152 2009c template into subjects’ T1w images in Structural Preprocessed for 7T package by the ANTs. The eccentricity maps of V1 were obtained in each individual’s native MNI space according to the structural retinotopic atlas by Benson, et al. (49), and the maps were warped into subjects’ T1w images in Structural Preprocessed for 7T package using the ANTs. The eccentricity maps of OR, we converted pRF results of each subjects’ OR into NIFTI files. Based on the eccentricity maps, we divided all ROIs into foveal (< 4 degree) and peripheral (> 4 degree) visual fields.

Effective connectivity of resting state fMRI data was analyzed using the SPM12. The full DCM model was consisted of feedforward (FF) and feedback (FB) connections between LGN and V1, as well as their self-connections (Figure 5A). The full DCM models for foveal and peripheral visual fields were analyzed and then averaged respectively by Bayesian fixed effect (FFX) averaging method (spm_dcm_average.m). The spectral DCM (sDCM) method was used (32), due to its applicability for resting state fMRI data (33). The group level analysis was performed using the Parametric Empirical Bayes (PEB) module of SPM (spm_dcm_peb.m). The commonality (group mean) and the eccentric OR activity (between subject effects) were specified as the two covariates of the design matrix. The retinotopy data (RETCW, RETCCW, RETEXP, RETCON, RETBAR1 and RETBAR2) were transformed into frequency domain by the fast Fourier transform (FFT). Then the eccentric OR neural activity was defined as the averaged magnitude at the fundamental stumuli frequency (∼0.031 Hz) (6) within the foveal and peripheral OR masks, respectively. Finally, the Bayesian model comparison, reduction and averaging (BMC, BMR and BMA) were performed (spm_dcm_peb_bmc.m) to test OR’s effects on the FF and FB connection between the LGN and V1. We defined reduced models as a FF model and a FB model. BMA was thresholded based on the free energy.

## Availability of Data and Code

Anonymized data and code to reproduce the results presented here are available at https://github.com/mynamewh/white-matter-functional-activity/tree/main/WM_pRF.

## Author contributions

X.W., P.Z., H.W., Y.Z. and B.Q. designed research; H.W., X.W., Y.W. and D.Z. analyzed data; P.Z., X.W. and H.W. wrote the paper.

## Acknowledgements

We thank Jia-Hong Gao and Sheng He for providing comments on an earlier version of the manuscript. This work was supported by the National Natural Science Foundation of China (grant nos. 81701665, 31871107, 31930053, 32070990, 21876041), Strategy Priority Research Program of Chinese Academy of Science (XDB32020200), Beijing Municiple Science and Technology Commission (Z181100001518002), State Key Laboratory of Ophthalmology, Optometry and Visual Science (325027).

