## Supplemental Figure 1 and 2 for "BOLD fMRI signals of visual white matter encode visuotopic information and predict effective connectivity between visual areas"

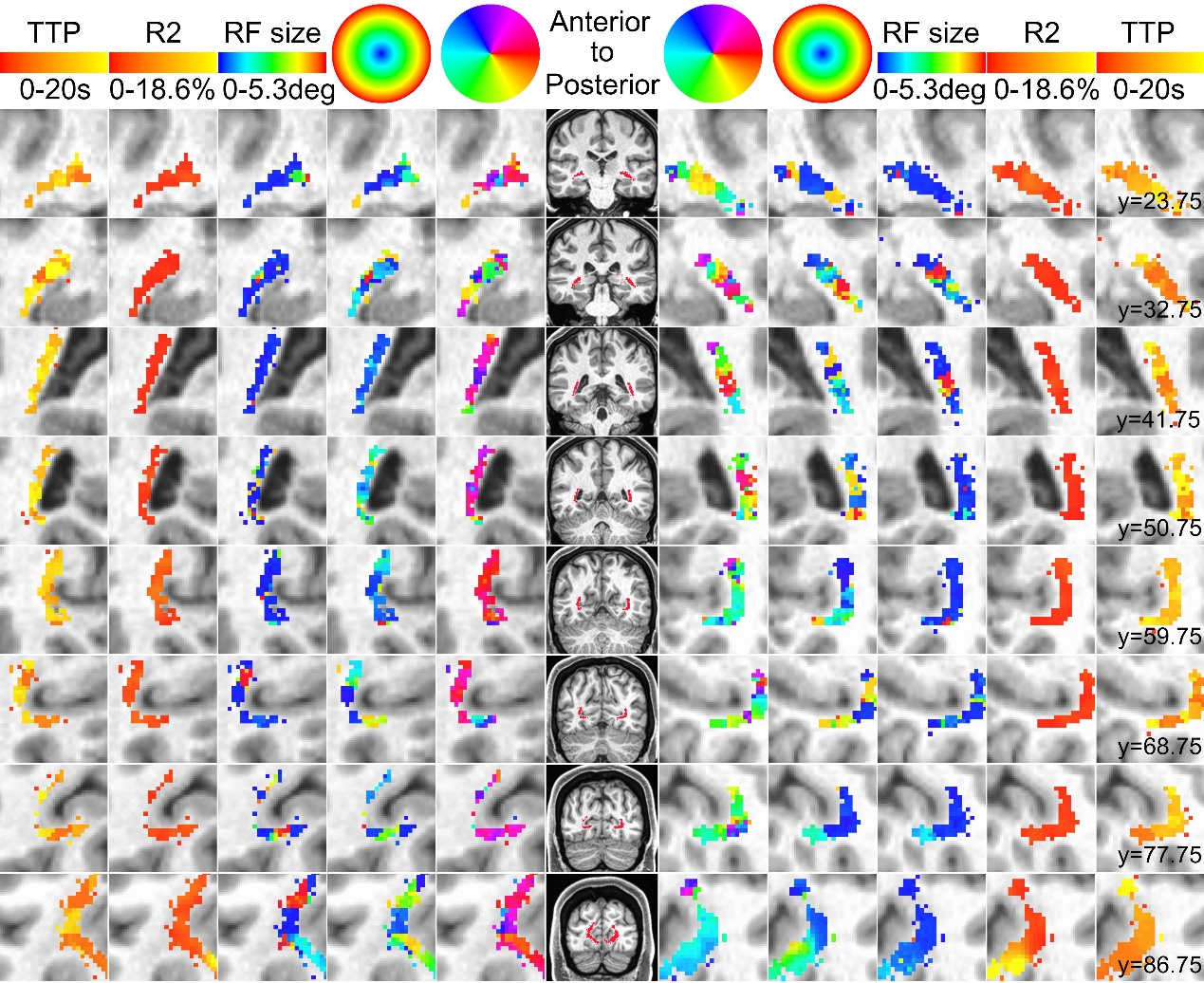

Figure S1. pRF maps and HRF delay for OR from a representative subject (HCP ID 144226). Related to Figure 2. All voxels within the OR mask are shown, conventions as in Figure 2. Retinotopy data were nonlinearly warped to ICBM152 2009c space and were spatially smoothed using a 4 mm smoothing kernel. The OR mask was obtained from automatic fiber tracking and constrained within the whole brain WM mask, then nonlinearly warped to ICBM152 2009c MNI space. At the individual level, polar angle maps show clear contralateral representations. The eccentricity maps show central visual field representation in the middle portion of OR and peripheral visual field representations from dorsal and ventral portions.

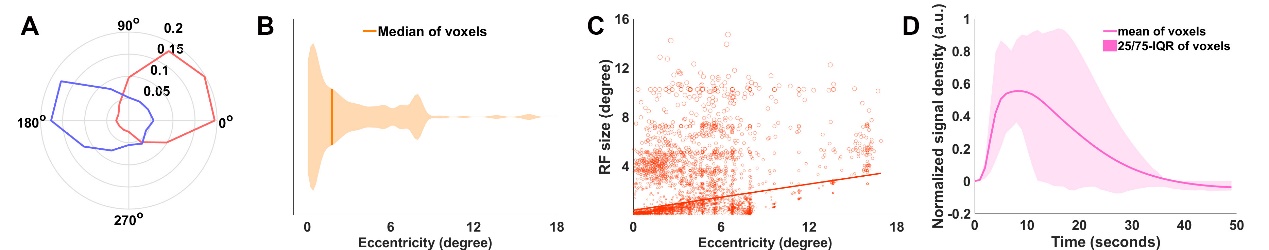

Figure S2. Model parameters of OR for the same subject. Related to Figure 4. (A) Polar angle distribution. (B) Eccentricity. 69.65% voxels represent 4 degrees of eccentricity in the central visual field. (C) The size of pRF increases with eccentricity (Pearson’s correlation, r = 0.32, p < 0.001). (D) Double gamma HRF model fitted HRF curve. The responses delay of HRF is 10.09 (mean) ± 5.17 (st.d.). Conventions as in Figure 4.
